# The landscape of intrinsic and evolved fluoroquinolone resistance in *Acinetobacter baumannii* includes suppression of drug-induced prophage replication

**DOI:** 10.1101/442681

**Authors:** Edward Geisinger, Germán Vargas-Cuebas, Nadav J. Mortman, Sapna Syal, Elizabeth L. Wainwright, David Lazinski, Stephen Wood, Zeyu Zhu, Jon Anthony, Tim van Opijnen, Ralph R. Isberg

## Abstract

The emergence of fluoroquinolone resistance in nosocomial pathogens has restricted the clinical efficacy of this antibiotic class. In *Acinetobacter baumannii*, the majority of clinical isolates now show high-level resistance due to mutations in *gyrA* (DNA gyrase) and *parC* (Topo IV). To investigate the molecular basis for fluoroquinolone resistance, an exhaustive mutation analysis was performed in both drug sensitive and resistant strains to identify loci that alter the sensitivity of the organism to ciprofloxacin. To this end, parallel fitness tests of over 60,000 unique insertion mutations were performed in strains with various alleles in genes encoding the drug targets. The spectrum of mutations that altered drug sensitivity was found to be similar in the drug sensitive and double mutant *gyrAparC* background having resistance alleles in both genes. In contrast, introduction of a single *gyrA* resistance allele, resulting in preferential poisoning of Topo IV by ciprofloxacin, led to extreme alterations in the insertion mutation fitness landscape. The distinguishing feature of preferential Topo IV poisoning was induction of DNA synthesis in the region of two endogenous prophages, which appeared to occur *in situ*. Induction of the selective DNA synthesis in the *gyrA* background was also linked to enhanced activation of SOS response and heightened transcription of prophage genes relative to that observed in either the WT or *gyrAparC* double mutants. Therefore, the accumulation of mutations that result in the stepwise evolution of high ciprofloxacin resistance is tightly connected to suppression of hyperactivation of the SOS response and endogenous prophage DNA synthesis.

**Importance:** Fluoroquinolones have been extremely successful antibiotics. Their clinical efficacy derives from the ability to target multiple bacterial enzymes critical to DNA replication, the topoisomerases DNA gyrase and Topo IV. Unfortunately, mutations lowering drug affinity for both enzymes are now widespread, rendering these drugs ineffective for many pathogens. To undermine this form of resistance, we sought to understand how bacteria with target alterations differentially cope with fluoroquinolone exposures. We studied this problem in the nosocomial pathogen *A. baumannii*, which causes resistant, life-threating infections. Employing genome-wide approaches, we uncovered numerous pathways that could be exploited to lower fluoroquinolone resistance independently of target alteration. Remarkably, fluoroquinolone targeting of Topo IV in specific mutants caused dramatic prophage hyperinduction, a response that was muted in strains with DNA gyrase as the primary target. This work demonstrates that resistance evolution via target modification can profoundly modulate the antibiotic stress response, revealing potential resistance-associated liabilities.

## Introduction

*Acinetobacter baumannii* is a frequent cause of multidrug resistant infections in hospitals and has been labeled a pathogen of critical priority for new drug development (1). This pathogen class has rapidly evolved a broad array of drug resistance mechanisms, limiting the usefulness of many widely-used antibiotics. A prime example is the fluoroquinolone class of antibiotics. These drugs are widely used to treat infections caused by a range of Gram-negative and Gram-positive bacteria, but they have been rendered obsolete against most *A. baumannii* isolates due to extremely high frequencies of resistance (2-4). Understanding how *A. baumannii* and related bacteria withstand treatment with fluoroquinolone antibiotics has the potential to lead to strategies to reverse or bypass resistance.

Fluoroquinolones inhibit DNA replication in bacteria by targeting two enzymes essential for DNA synthesis, the type II topoisomerases DNA gyrase (*gyrAB* genes) and topoisomerase IV (Topo IV, *parCE*). These enzymes modulate DNA topology to maintain negative DNA supercoiling (DNA gyrase) or decatenate newly replicated DNA (Topo IV), and in so doing, break and religate DNA. Binding of fluoroquinolones to these enzymes traps them in an intermediate state that is bound to cleaved DNA, resulting in double-strand DNA breaks, blocked replication fork progression and, at high drug concentrations, cell death (5).

Acquired resistance to fluoroquinolones commonly arises through stepwise mutations that disrupt the ability of the drug to bind its preferred target enzymes. In Gram-negative bacteria including *A. baumannii*, these mutations typically arise first in *gyrA*, encoding the GyrA subunit of DNA gyrase which is the more sensitive of the two enzyme targets (6). In the presence of resistant GyrA, the less sensitive Topo IV (encoded by *parC*) becomes the target and the site of second-step resistance mutations. In addition to target site alterations, acquisition of mutations that upregulate drug efflux pumps or accessory genes that allow drug modification enable bacteria to develop fluoroquinolone resistance (7). A large fraction of *A. baumannii* isolates harbor target-site mutations in *gyrA* and *parC* (8, 9) and mutations causing overproduction of one or more RND-class efflux systems that act on fluoroquinolone drugs (AdeABC, AdeFGH, AdeIJK (10-12)).

Acquired resistance mechanisms generally act in combination with intrinsic resistance strategies in a cumulative manner to raise the amount of fluoroquinolone antibiotic required to block bacterial growth. Regulated production of native efflux pumps contributes to intrinsic fluoroquinolone resistance in many bacteria (13). Of the RND systems in *A. baumannii*, native levels of AdeIJK in wild-type (WT) strains lacking acquired mutations have been shown to provide intrinsic resistance to fluoroquinolones (12). Whether regulated production of other efflux systems provides intrinsic fluoroquinolone resistance is less clear.

Another major strategy for intrinsic fluoroquinolone resistance is activation of DNA damage repair pathways (14). DNA lesions caused by fluoroquinolone intoxication are processed to single-stranded DNA and subsequently induce the SOS repair response, resulting in de-repression of many genes involved in DNA recombination and repair (15). Knockout mutations in a variety of DNA repair genes result in increased fluoroquinolone susceptibility in several species (14-24). In certain cases, the SOS response also induces mobile genetic elements that carry antibiotic resistance or toxin genes, potentially influencing the spread of resistance or virulence traits (25). The *A. baumannii* SOS repair response is non-canonical, lacking clear orthologs of many major players in other systems (26) and is characterized by a phenotypically variable response within cell populations (27). Inactivation of RecA, a central protein mediating DNA recombinational repair and SOS induction, or the RecBCD Exonuclease V complex responsible for double-strand break repair, greatly raises fluoroquinolone sensitivity in *A. baumannii* (28, 29). The role of the SOS response and other DNA repair systems, however, in the development of antibiotic resistance in this organism is largely unknown.

In this study, we present the results of a comprehensive screen for determinants of intrinsic resistance to the fluoroquinolone antibiotic ciprofloxacin in *A. baumannii*. We hypothesized that the ciprofloxacin resistome varies depending on the drug target (DNA gyrase or Topo IV) that is preferentially poisoned, which is determined by possession of WT or resistant versions of the enzymes. We therefore performed parallel screens with isogenic *A. baumannii* strains containing sensitive or resistant *gyrA* and *parC* alleles to uncover the influence of target selectivity on the intrinsic resistance landscape. This analysis led to the surprising discovery that endogenous prophage activation by fluoroquinolones shows dramatic dependence on the availability of a sensitive *parC* allele.

## Results

### Identification of *Acinetobacter baumannii* loci that confer altered sensitivity to ciprofloxacin

As part of a largescale effort to characterize the molecular nature of intrinsic resistance of *Acinetobacter baumannii* to antimicrobials, we identified the entire spectrum of insertion mutations that cause altered sensitivity to the fluoroquinolone antibiotic ciprofloxacin during growth in bacteriological culture. A number of studies have demonstrated that mutations that cause antibiotic hypersensitization in strain backgrounds lacking demonstrable resistance loci exhibit these effects independently of whether there are target site resistance mutations or antibiotic-inactivating enzymes present in the strains being interrogated (20, 30). We wanted to test this model by first identifying loci that confer intrinsic resistance in a strain background having intact drug targets, and then comparing them with intrinsic resistance loci identified in strains having drug target mutations in DNA gyrase or Topo IV. For this work, we will refer to lesions resulting in lowered resistance to the antibiotic as ciprofloxacin-hypersensitizing mutations in each strain background, and the genes harboring these mutations as loci of hypersensitization.

To identify ciprofloxacin hypersensitization loci, ciprofloxacin concentrations below the minimal inhibitory concentration (MIC) (Table 1) were identified that resulted in growth rates of *A. baumannii* ATCC17978 in rich broth that were between 60-80% of that observed without antibiotics (Fig. 1A). Three of these concentrations were chosen for further analysis (0.05, 0.075 and 0.09-0.10 μg/ml) to determine the relative fitness of insertion mutations when subjected to each of the antibiotic stress conditions. Multiple independent Tn*10* insertion pools (7 pools for 0.05 μg/ml, and 11 pools for 0.075 and 0.09-0.10 μg/ml ciprofloxacin) having between 6,000 and 18,000 individual insertions (60,000 separate sites in all) were grown in broth in the presence or absence of ciprofloxacin for approximately 8 generations. DNA samples taken from the initial time point prior to growth (t_1_) and the final timepoint after 8 generations growth (t_2_) were prepared from each of the pools. The insertion sites were then amplified preferentially and subjected to high density sequencing, followed by determining the relative fitness of each insertion mutant based on density of reads (Materials and Methods; Data Set S1). Using accepted strategies, the fitness of each insertion mutant strain was calculated relative to the entire pool (31). To standardized results across experiments, fitness values were normalized to insertions found in 18 neutral sites located in pseudogenes or endogenous transposon-related genes throughout the genome (“neutral” mutants), to allow an accurate quantitation of the representation of mutants relative to control insertions predicted to have no effect on growth (32). The normalized data from the individual insertion mutations were aggregated for each gene to calculate a mean fitness level for the entire spectrum of mutations found within a particular gene. The complete datasets were then displayed on an individual gene level as the growth rate changes for mutations relative to the growth rate of the entire pool (Fig. 1B). Candidate mutants were identified that showed lower (hypersensitizing loci) or higher fitness levels based on the criteria that the False Discovery Rate (FDR) q value was <0.05, a change in fitness (W_diff_) was >10%, and fitness value was derived from at least 3 independent insertion mutants ((33); Materials and Methods).

**Table 1.** Minimal Inhibitory Concentration (μg/ml) of Ciprofloxacin with WT and mutant *A. baumannii.*

| strain | amino acid change: |  | ciprofloxacin<br>MIC (µg/ml) |
| --- | --- | --- | --- |
|  | GyrA | ParC |  |
| WT | - | - | 0.25 |
| *gyrA*<sup>R</sup> | S81L | - | 2 |
| *parC*<sup>R</sup> | - | S84L | 0.25 |
| *gyrA*<sup>R</sup> *parC*<sup>R</sup> | S81L | S84L | 32 |

**Fig. 1.**
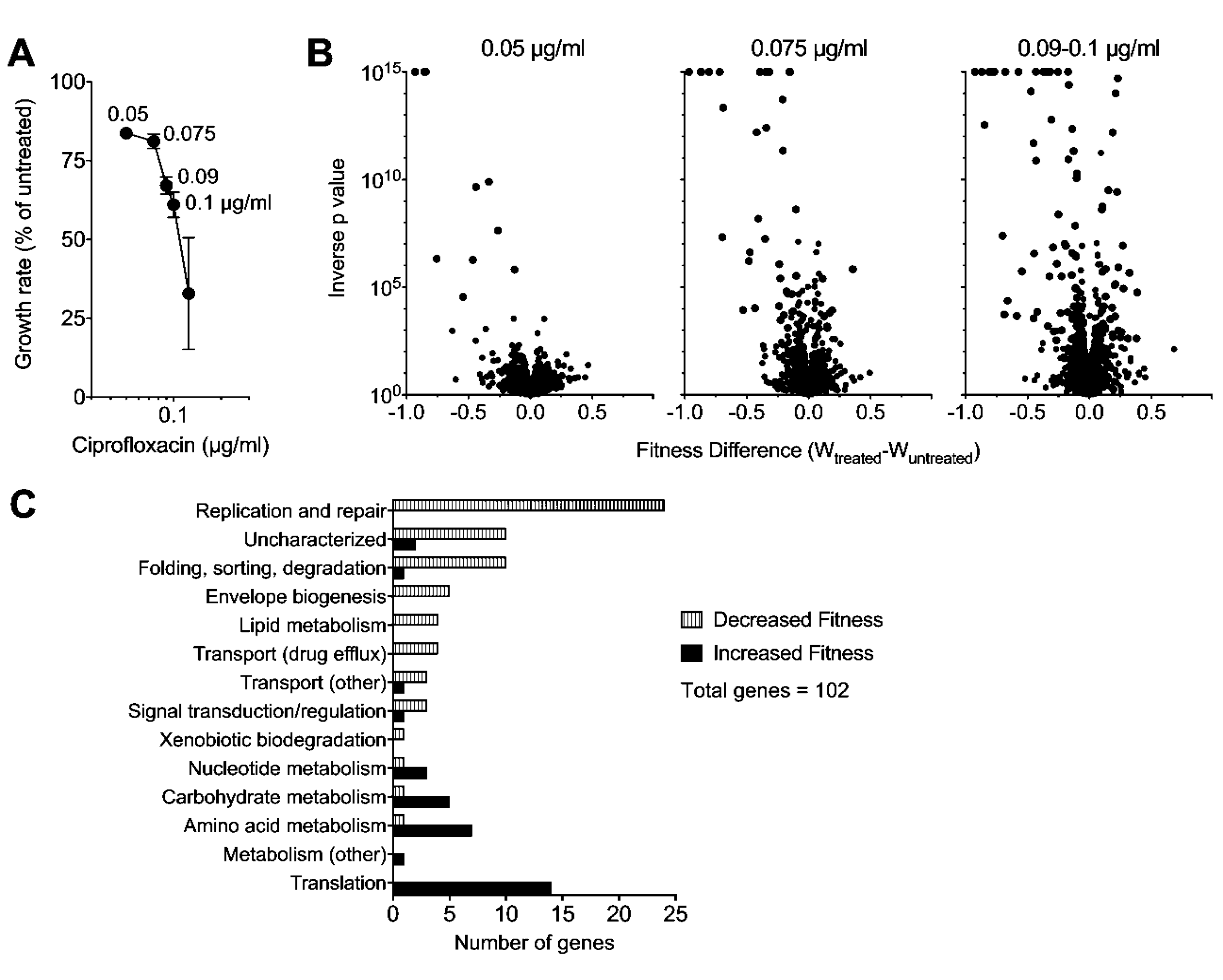
Tn-seq quantification of genome-wide mutant fitness in *A. baumannii* (*gyrA*^WT^ and *parC*^WT^) during growth with sub-MIC ciprofloxacin. (A) Graded concentrations of ciprofloxacin cause increasing degrees of growth inhibition. Transposon mutant libraries constructed in a strain background harboring WT alleles of the *gyrA* and *parC* genes were grown with ciprofloxacin at the concentrations indicated. Growth rate relative to untreated control was determined from bacterial density measurements. Data points show average ± SD (n ≥ 2). (B) Tn-seq fitness profiles during ciprofloxacin challenge. Mutant pools were grown with or without the indicated ciprofloxacin concentration and average fitness for each chromosomal gene was calculated. Change in fitness resulting from ciprofloxacin treatment relative to untreated controls is plotted against significance score resulting from parallel t-tests. Data points shaded in black indicate gene knockouts causing significant alteration in fitness during drug challenge (W_treated_-W_untreated_ had a magnitude > 0.1 and FDR <0.05). (C) Functional categories of significant gene hits determining fitness during challenge with 0.09-0.1 μg/ml ciprofloxacin. Information from KEGG and UniProt functional annotations and from orthologs in well-studied reference species were used to place genes into the listed broad categories.

Small increases in the dose of ciprofloxacin greatly increased the spectrum of ciprofloxacin hypersensitivity loci (Fig. 1B; black). At a dose causing approximately 20% growth inhibition, mutations in only 10 genes passed the criteria for lower fitness relative to the rest of the pool in the presence of the drug. These included insertions in: two genes involved in double strand break repair (*recB* and *ruvA*, encoding subunits of exonuclease V and the Holiday junction helicase); the major egress pump which is often found overproduced in clinical strains having high level fluoroquinolone resistance (*adeIJK*); and *ctpA*, a periplasmic protease shown to be a target of mutations that augment β-lactam resistance in strains lacking the *bfmRS* global regulatory system (34) (Data Set S1). Increasing the drug dose had two effects on expanding the spectrum of hypersensitivity loci. First, although the number of hypersensitivity loci that contribute to the enzymology of DNA repair increased from 2 members to 20 in the high dose regimen, this expansion largely involved hitting additional subunits of the same complexes or backup systems of the enzymes identified in the low dose regimen (*recBCD, sbcCD*, *ruvABC*) (Data Set S1). This emphasizes the importance of protecting against double strand breaks caused by fluoroquinolone-poisoned DNA gyrase (35). Secondly, increasing dose resulted in hypersensitivity loci in cell envelope integrity proteins, additional protein-processing enzymes, and a MATE class proton-driven efflux pump (*abeM*) shown to export ciprofloxacin and other antibiotic compounds when cloned in *E. coli* (36) (Data Set S1; Fig. 1C). Interestingly, increasing the drug dose did not implicate the two other major RND efflux systems in protecting from ciprofloxacin stress even though they are known to provide low-level resistance after overproduction (12). This may be explained by the fact that the *adeIJK* system is the only RND egress pump known to have a high basal level of expression in WT strains lacking acquired resistance mutations, while the inducing signals for the other systems have not been identified (12). Strikingly, at the higher drug doses, mutations in *adeN*, which encodes the negative regulator of *adeIJK,* increased the fitness of *A. baumannii* relative to the rest of the pool (Data Set S1). These data argue strongly that the primary efflux pumps involved in intrinsic protection from fluoroquinolone stress are AdeIJK and AbeM.

In addition to hypersensitivity loci, mutations were identified at the highest drug dose that resulted in increased fitness relative to the insertion pool (Fig. 1B and C, Data Set S1). The mutations that most frequently increased fitness targeted nonessential components of the protein translation machinery, particularly enzymes that post-translationally modify tRNA, rRNA and assembly of ribosomal protein complexes. That disruption of this circuit is tightly associated with increased drug resistance is consistent with studies showing that a spectrum of antibiotic resistant isolates in different species evolve mutations causing slowed translation rate (37, 38). The results are also consistent with a study demonstrating that lowering ribosomal synthesis increases resistance to ciprofloxacin by restoring an optimal balance between protein and DNA synthesis levels during DNA stress (39)). Most notable among the insertions identified were those in *gidA*, which is part of a complex involved in 5-methylaminomethyl-2-thiouridine (mnm^5^s^2^U34) modification of tRNAs (Data Set S1; (40)). We have previously identified this gene as an additional target of mutations bypassing drug hypersensitivity resulting from loss of *bfmRS* (34), indicating the tight connection between mutations in this gene and drug resistance.

### Deletion mutants have drug sensitivities predicted by Tn-seq

Targeted deletion or null mutations were isolated in nonessential genes predicted to have altered drug sensitivity in the presence of ciprofloxacin. The mutations were chosen based on their fitness in the Tn-seq analysis, the magnitude of the effects predicted, and differing functional categories (Fig. 1C). For instance, mutations in the egress pump-encoding *adeIJK* showed extremely poor fitness and were rarely recovered after growth in ciprofloxacin (Fig. 2A). Similarly, the insertions in *ctpA* showed very low fitness. In contrast, although ciprofloxacin treatment lowered fitness for mutants lacking the penicillin binding protein PBP1A, these mutations clearly had weaker effects in the Tn-seq analysis. When this set of targeted mutants was analyzed further, loss of *adeIJK, recN, ctpA* and *pbp1A* all resulted in heightened drug sensitivity (Fig. 2B). In contrast, deletion of *gidA-*encoded tRNA modification enzyme resulted in enhanced fitness in the presence of ciprofloxacin, with increased yields in broth cultures exposed to 0.15 μg/ml of antibiotic (Fig. 2B).

**Fig. 2.**
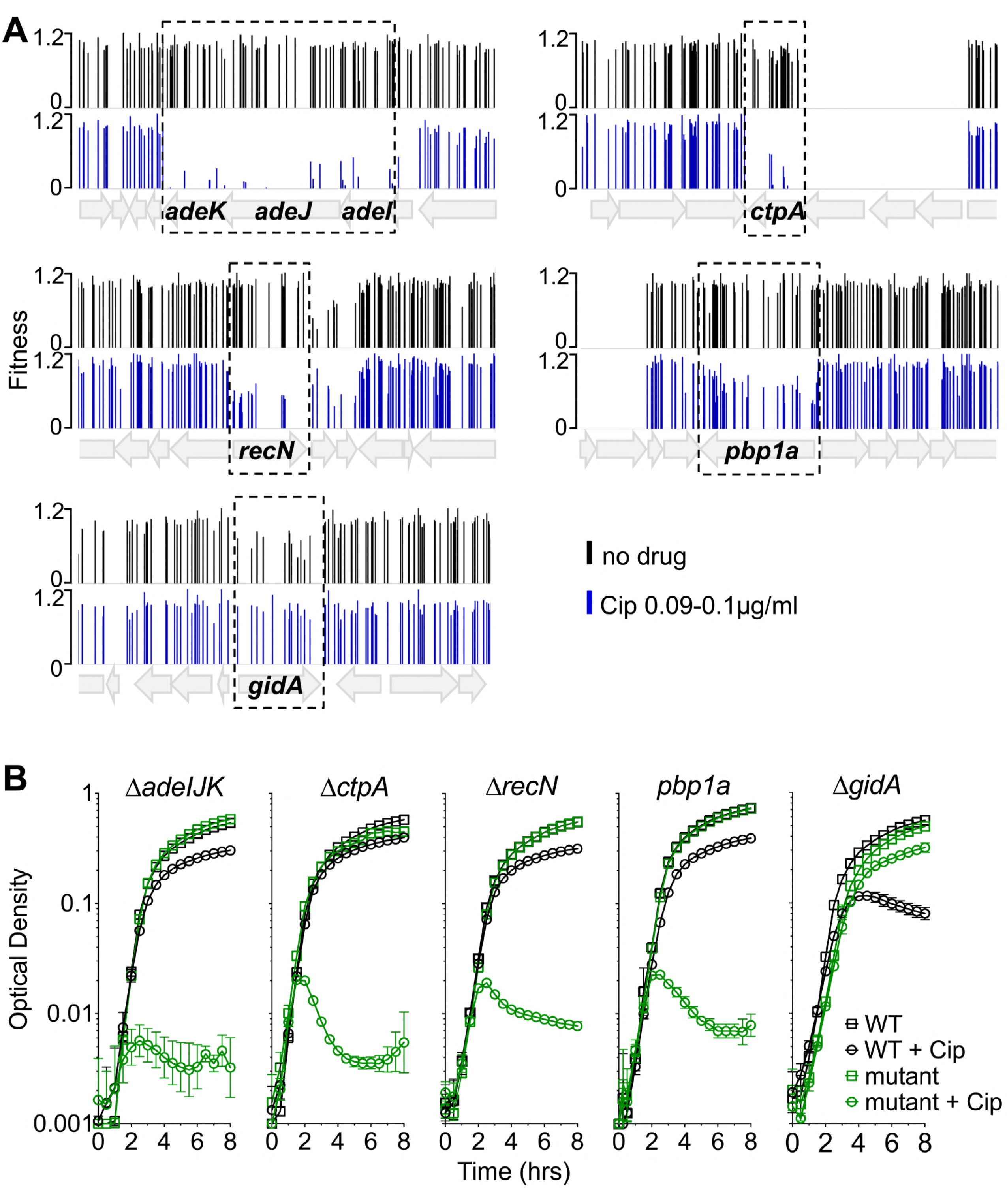
Magnitude of growth impairment is predicted by the severity of the Tn-seq fitness defect. (A) Tn-seq fitness profiles of transposon mutant pools constructed in a *gyrA*^WT^*parC*^WT^ strain. Pools were grown without or with ciprofloxacin at 0.09-0.1 μg/ml. Bars show fitness values of each transposon mutant at the indicated locus across all tested pools. (B) Growth of pure cultures of WT or the indicated mutant in the absence or presence of ciprofloxacin (0.09 μg/ml for ∆*ctpA* and *pbp1A*, 0.1 μg/ml for ∆*adeIJK* and *∆recN*, and 0.15 μg/ml for ∆*gidA*). The *pbp1a* mutant tested was *pbp1a*(N178TfsX27). Data points show geometric mean ± SD (n = 3). Cip, ciprofloxacin.

### Identification of loci that result in altered ciprofloxacin sensitivity in *A. baumannii* target site mutants

A majority of the current clinical isolates of *A. baumannii* are resistant to fluoroquinolones, and these isolates commonly have the *gyrA*(S81L) and *parC*(S84L) target site mutations that lower the affinity for these antibiotics (41). To determine the spectrum of insertions that cause altered sensitivity to ciprofloxacin in strains having resistance alleles, *gyrA*(S81L) (hereafter referred to as *gyrA*^R^) and *gyrA*(S81L) *parC*(S84L) (referred to as *gyrA*^R^ *parC*^R^) mutants were generated, and each strain was subjected to Tn*10* mutagenesis. Pools totaling more than 70,000 insertion mutations were constructed in each background. Insertion pools were challenged with ciprofloxacin, using drug concentrations below the MIC (Table 1) that resulted in 30-40% growth inhibition for each strain (1.1μg/ml for *gyrA*^R^; 13-14 μg/ml for *gyrA*^R^ *parC*^R^ double mutant; Fig. 3A). The spectrum of insertions that resulted in hypersensitivity to ciprofloxacin in the *gyrA*^R^ *parC*^R^ double mutant strain backgrounds was very similar to the WT (Fig. 3C,D). In fact, almost every ciprofloxacin hypersensitive locus in the double mutant background was identified previously in the WT (green circles, Fig. 3C; Data Sets S1 and S3). In addition, there was a number of hypersensitivity loci identified in the WT pools that did not pass the discovery criteria in the double mutant (FDR<0.05; W_diff_>0.1). A number of these below-threshold candidates in the *gyrA*^R^ *parC*^R^ double mutant strain background encoded subunits of the proteins identified as ciprofloxacin-hypersensitive loci (green circles, Fig. 3C; Data Set S3). These results are similar to what we had observed in our graded series of drug treatments of insertion pools in the WT strain, indicating that the results from the WT strain and the drug resistant double mutant are largely the same.

**Fig. 3.**
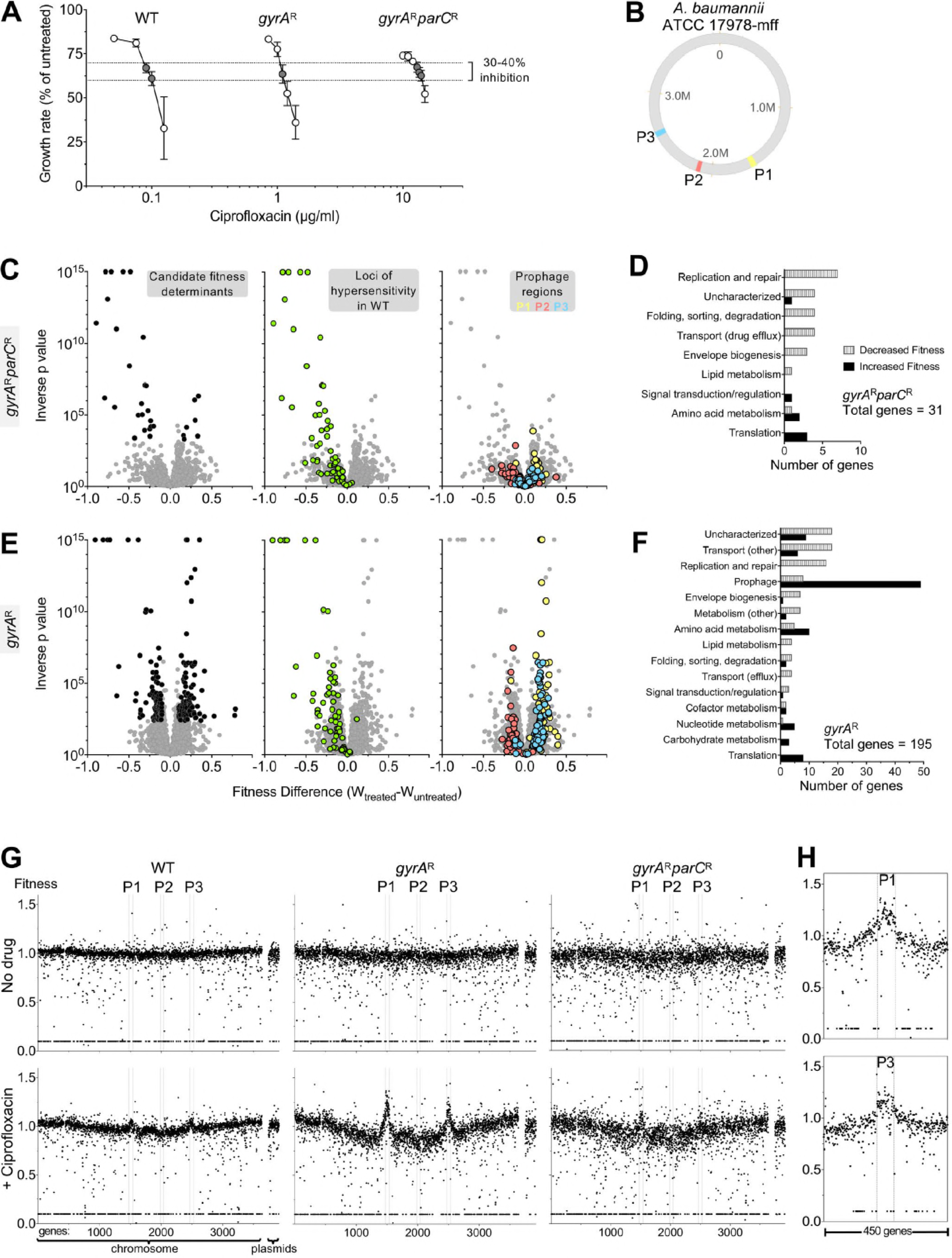
Acquisition of *gyrA* resistance allele dramatically alters *A. baumannii* Tn-seq profile during ciprofloxacin challenge. (A) Transposon pools constructed in strains harboring resistance alleles in *gyrA* or both *gyrA* and *parC* require increasing concentrations of ciprofloxacin to result in growth inhibition. Growth rate inhibition relative to untreated pools was plotted as in Fig. 1A. *gyrA*^WT^ *parC*^WT^ growth data are from identical experiment shown in Fig. 1A and are displayed to allow comparison to behavior of drug resistant mutants. Data points show average ± SD (n ≥ 2). Samples from cultures with 30-40% growth inhibition (dotted lines) were processed for Tn-seq. (B) Location of prophage regions (P1-P3) on *A. baumannii* 17978-mff chromosome map. Prophage positions were identified by using the PHASTER database (66). (C-F) *gyrA* resistance allele influences Tn-seq fitness profiles associated with ciprofloxacin stress. Mutant pools were challenged with drug concentrations that resulted in equivalent 30-40% growth inhibition [*gyrA*^R^, 1.1 μg/ml; *gyrA* ^R^*parC* ^R^, 13-14 μg/ml]. (C, E) Tn-seq fitness scores for each chromosomal gene with the indicated strain were calculated and visualized as in Fig. 1B (leftmost subpanel). Middle and rightmost subpanels show the identical dataset, with highlighting of loci for which knockout causes ciprofloxacin hypersensitization in the WT genetic background (green), or loci within prophages (color indicated in key). (D, F) Gene hits associated with significant changes in fitness during treatment were placed into functional categories as in Fig. 1C. Tn-seq hits resulting from *gyrA*^R^ libraries treated with ciprofloxacin are enriched in prophage genes (F). (G) Tn-seq fitness scores resulting from ciprofloxacin challenge show genome positional bias that is greatly amplified in *gyrA*^R^ mutant pools. Average per-gene Tn-seq fitness values are plotted in order of gene position on the chromosome or on plasmids pAB1-3. Boundaries of prophage regions (P1-P3) are indicated by vertical dotted lines. Top, no drug control. Bottom, ciprofloxacin was added at the concentrations indicated in panel A resulting in 30-40% inhibition (WT, 0.09-0.1 μg/ml; *gyrA*^R^, 1.1 μg/ml; *gyrA*^R^*parC*^R^, 13-14 μg/ml). (H) Expanded view of per-gene Tn-seq fitness scores in regions surrounding prophages P1 and P3 for *gyrA*^R^ mutant treated with 1.1 μg/ml ciprofloxacin.

The results from the *gyrA*^R^ single mutant background, however, diverged greatly from the WT and the *gyrA*^R^ *parC*^R^ double mutant (Fig. 3E). A large fraction of insertions were identified that altered drug sensitivity to ciprofloxacin, with a surprising number showing increased fitness during drug exposure (Fig. 3E). Over 40 of the insertions that exhibited increased fitness were located in putative prophage genes (blue and yellow circles, Fig. 3E; Fig. 3F) from two of the three predicted phages integrated into the bacterial chromosome (Fig. 3B; Data Set S2). No such fitness changes were seen in the WT (Fig. S2, Data Set S1) or *gyrA*^R^ *parC*^R^ double mutant (Fig. 3C, Data Set S3). To analyze this result further, the normalized fitness of mutations in each gene was plotted as a function of position on the chromosome. In the absence of antibiotic, there was no clear positional effect of altered fitness levels along the length of the chromosome (Fig. 3G). In contrast, in the presence of antibiotic, there was an apparent increase in fitness levels centered within chromosomal locations harboring prophages P1 and P3 in the *gyrA*^R^ single mutant (Fig. 3G). Although some of this effect could be explained by loss of prophage gene function resulting in enhanced fitness, insertions in chromosomal regions near, but outside, the prophage boundaries similarly showed apparent increases in fitness relative to the rest of the chromosomal insertions (Figs. 3G and H). As fitness levels are measured by counting the number of reads in specific regions of DNA, this phenomenon is consistent with selective local amplification of chromosomal material that initiates within these prophages, extending outward from the integration sites into nearby DNA regions.

### Two prophage regions are selectively amplified in response to ciprofloxacin in the single *gyrA*^R^ mutant

We next tested the model that there is induction of DNA synthesis in the region surrounding two of the chromosomally-located prophage clusters. Purified single colonies from the WT, *gyrA*^R^, and *gyrA*^R^*parC*^R^ double mutant strains were grown in broth culture for 3.5 hours in the presence of four different concentrations of ciprofloxacin that ranged from 30-80% growth inhibition and compared to bacteria grown in the absence of drug (Fig. 4A). DNA was then prepared from each of the cultures and subjected to whole genome sequencing using an average read length of 100 bp. The density of these individual short reads was plotted as a function of the chromosomal coordinates, to identify regions of chromosomal DNA that were selectively amplified in the presence of drug (Fig. 4B). Analysis of the *gyrA*^R^ single mutant showed hyperamplification of prophages 1 and 3, with read density in the prophage regions observed as a function of drug concentration. In contrast, there was little evidence of this selective amplification in the WT strain, while the *gyrA*^R^*parC*^R^ largely reversed these effects. Consistent with the Tn-seq data, there was amplification of DNA extending beyond the prophage-chromosomal DNA junction, indicating that drug-driven DNA synthesis was initiated *in situ* and continued beyond the ends of the prophages into adjacent chromosomal DNA (Fig. 4C). We conclude that in a *gyrA^R^* background, selective blockage of the *parC*-encoded Topo IV protein resulted in DNA synthesis induction in these two prophage regions.

**Fig. 4.**
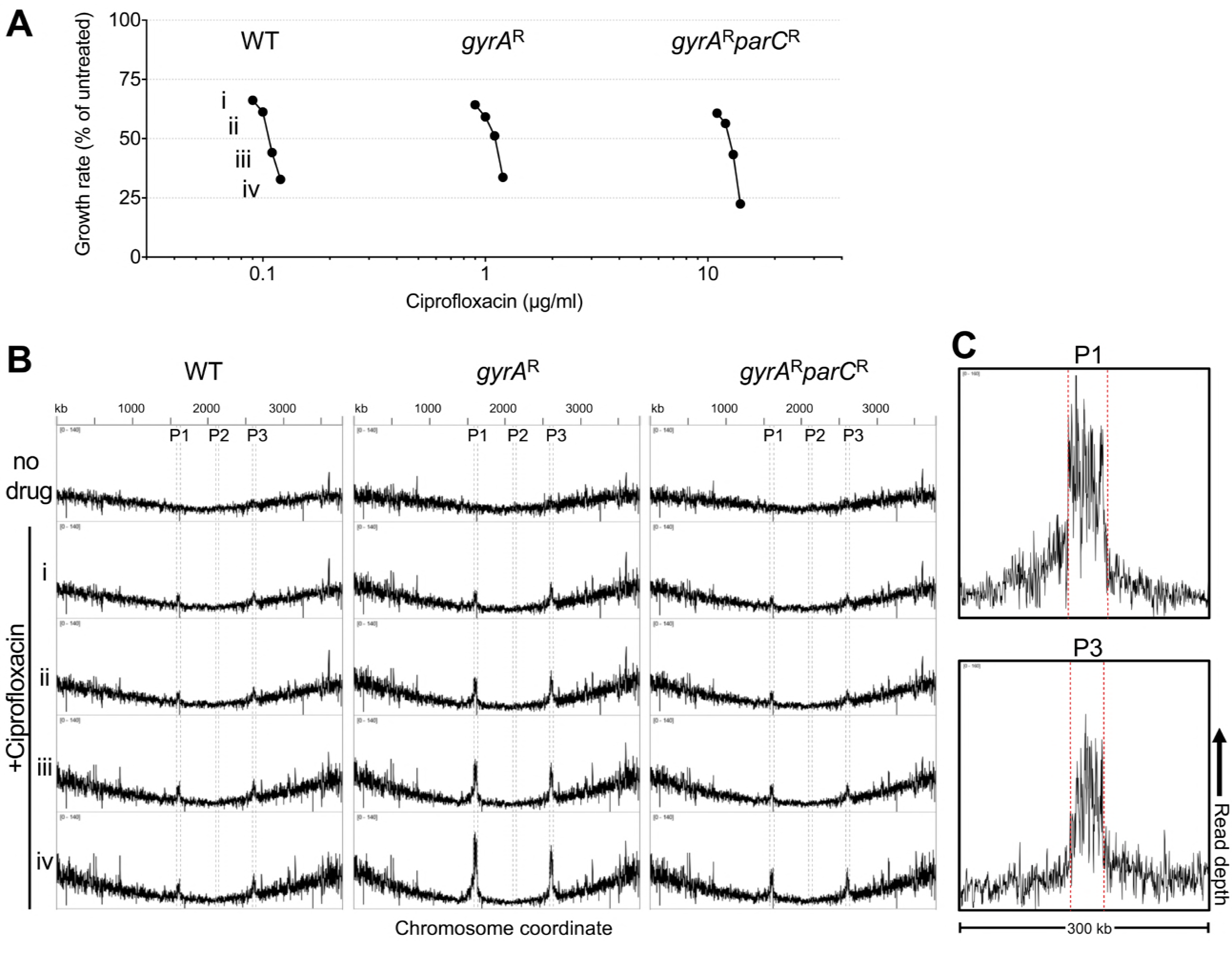
Ciprofloxacin-induced amplification of prophage DNA in strains harboring the *gyrA*^R^ single-step resistance genotype. (A) Pure cultures of strains of the indicated genotype were challenged with graded ciprofloxacin doses resulting in four levels of growth inhibition (Roman numerals). (B) DNA content from each culture was analyzed by deep sequencing. x-axis indicates nucleotide position along the *A. baumannii* ATCC17978-mff chromosome. y-axis indicates normalized read depth (0-140 counts per million). Boundaries of prophage regions P1-3 are indicated in red. Roman numerals indicate the level of growth inhibition caused by ciprofloxacin. Data are representative of two independent experiments. (C) Expanded view of 300kb window showing amplification of genomic regions surrounding prophages P1 and P3. Y-axis indicates normalized sequencing read depth (0-160 counts per million).

To determine if transcription of prophage genes is specifically amplified in the *gyrA*^R^ mutant relative to the WT, the two strains were grown in triplicate cultures in two concentrations of antibiotic for 3.5 hours that gave between 40-70% growth inhibition over approximately 7 generations (Fig. 5A). The cells were then extracted, subjected to RNAtag-seq analysis (42), and the ratio of transcription for each gene in the presence/absence of ciprofloxacin was displayed as a function of chromosomal map position (Fig. 5B, Data Set S4). There was preferential amplification of transcription of prophage genes in the presence of antibiotic treatment in both strain backgrounds (Fig. 5B). Furthermore, transcription was hyperactivated in all three prophages, including prophage 2 which showed no evidence of preferential DNA amplification (compare Figs 4B and 5B). Higher expression levels were observed with prophage genes in the *gyrA*^R^ single mutant compared to WT (Fig. 5B), and these levels were also apparent when directly comparing transcription in WT and *gyrA*^R^ after ciprofloxacin treatment (Fig. S3). In *gyrA*^R^-single mutants, enhanced expression in response to ciprofloxacin extended beyond the prophage-chromosomal DNA junctions with prophage 1 and to some extent with prophage 3 (Fig. 5C and S3), consistent with increased DNA template availability partially contributing to heightened transcription in this strain background. Hyperexpression in response to ciprofloxacin terminated at the prophage-chromosomal DNA junctions with prophage 2, consistent with the observation that this region experienced no DNA amplification (Fig. 5C). These results indicate that preferential blockage of Topo IV in the single mutant results in hyperactivation of prophage transcripts.

**Fig. 5.**
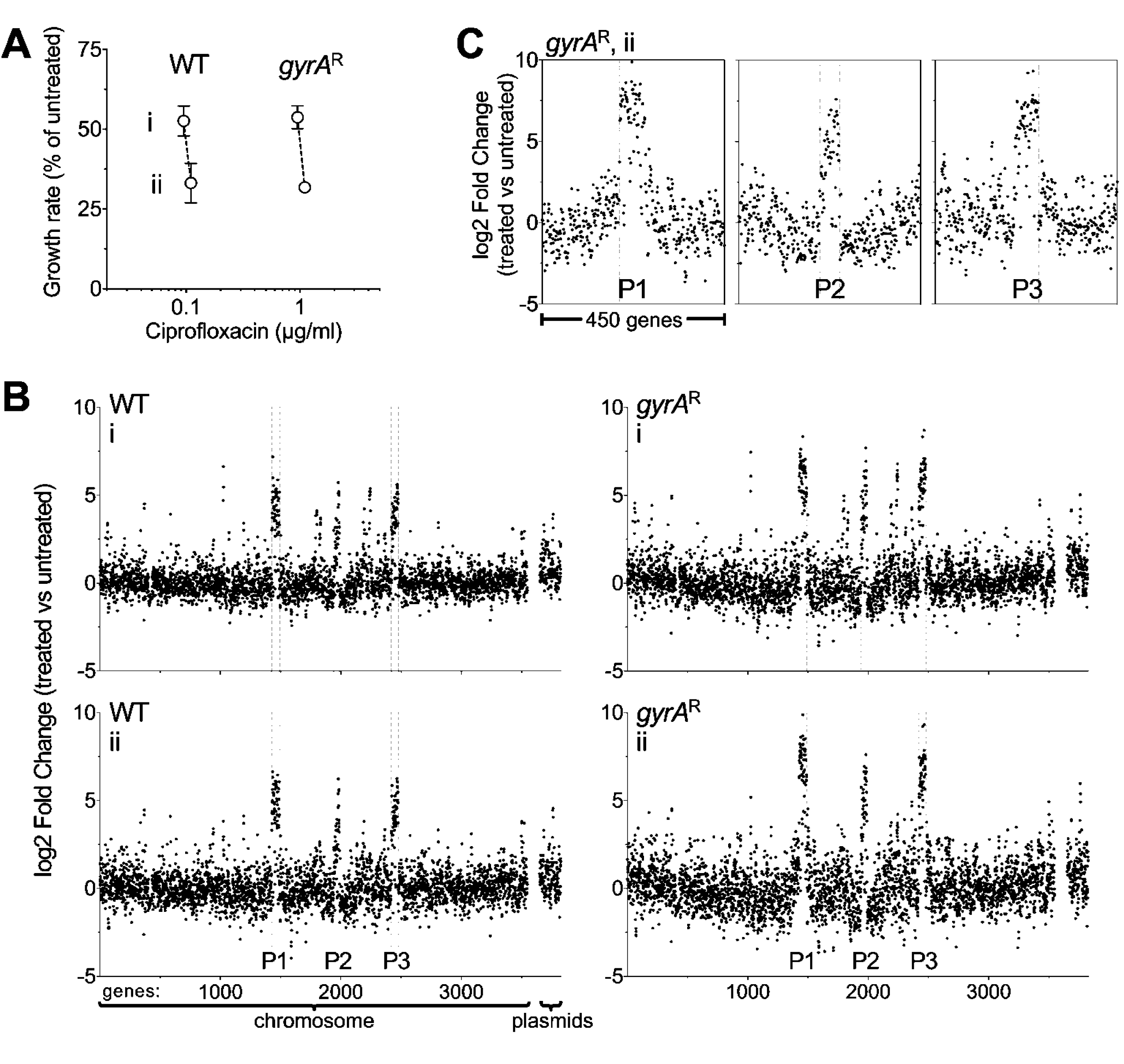
Ciprofloxacin challenge results in activation of prophage gene expression that is heightened in *gyrA*^R^-single mutants. (A) Strains of the indicated genotype were challenged with ciprofloxacin concentrations that resulted in two levels of growth inhibition relative to no treatment (i, ~45% growth inhibition; ii, ~70% growth inhibition). Data points show average ± SD (n = 3). (B). RNA-seq transcriptional profiles of panel A cultures. Fold change (log2) of each gene (treated vs untreated) was plotted in order of position on the chromosome or plasmids (pAB1-3). rRNA and tRNA genes were excluded from RNA-seq analysis, resulting in different gene number assignments as compared to those in Fig. 3D. Boundaries of prophage regions P1-3 are denoted by vertical dotted lines. Roman numerals indicate growth inhibition level. (C) Expanded views of RNA-seq log2-fold change ratios for genes surrounding P1-3 in *gyrA*^R^-single mutant (condition *ii*, ciprofloxacin 1.1μg/ml).

Intoxication of bacterial topoisomerase enzymes by fluoroquinolone antibiotics induces DNA damage, driving an SOS response (16, 43). We investigated the extent to which ciprofloxacin-induced hyperactivation of prophage gene expression coincided with SOS response induction, and whether *gyrA* or *parC* resistance alleles influenced this response. Several genes associated with the SOS response (27, 43, 44) showed heightened transcription as a consequence of ciprofloxacin treatment (Fig. 6A). For several SOS genes, transcript induction was significantly higher in *gyrA*^R^ compared to WT (Fig. 6A; asterisks). These included genes adjacent to prophage-chromosomal DNA junctions (*umuC* and *umuD* paralogs) as well as those not directly linked to prophages (*recA*, *gst*; Fig. 6B). RecA is a key component of the SOS response that is induced by DNA damage in *A. baumannii* and is critical for withstanding ciprofloxacin stress independent of the background resistance genotype (see Data Sets S1-S3). To analyze the interplay of SOS induction by ciprofloxacin with target availability at the level of single cells, we utilized a plasmid-based transcriptional fusion of the *recA* regulatory elements (promoter and 5’-untranslated region) to the fluorescent reporter *mKate2* (45). WT, *gyrA*^R^, and *gyrA*^R^*parC*^R^ strains harboring the reporter fusion were cultured in the presence of graded levels of ciprofloxacin, and reporter signal was measured in individual cells by fluorescence microscopy (Materials and Methods). Increasing sub-MIC doses of ciprofloxacin caused increasing degrees of induction of the *recA* reporter in all strain backgrounds (Fig. 6C). Notably, reporter activity was approximately 2-fold higher in the *gyrA*^R^ single mutant than in the WT or double mutant at equivalent levels of growth inhibition (Fig. 6C). Varying degrees of *recA* induction within populations of *gyrA*^R^ single mutant cells were observed, and this variability roughly matched that observed with WT (Fig. S4). Increased signal in the *gyrA*^R^ strain was not observed with a control reporter fusion to a gene that is nonresponsive to ciprofloxacin (*trpB*p-UTR) ((45), Fig 6D), indicating that the SOS transcriptional response was specifically enhanced as a consequence of ciprofloxacin inhibition of Topo IV.

**Fig. 6.**
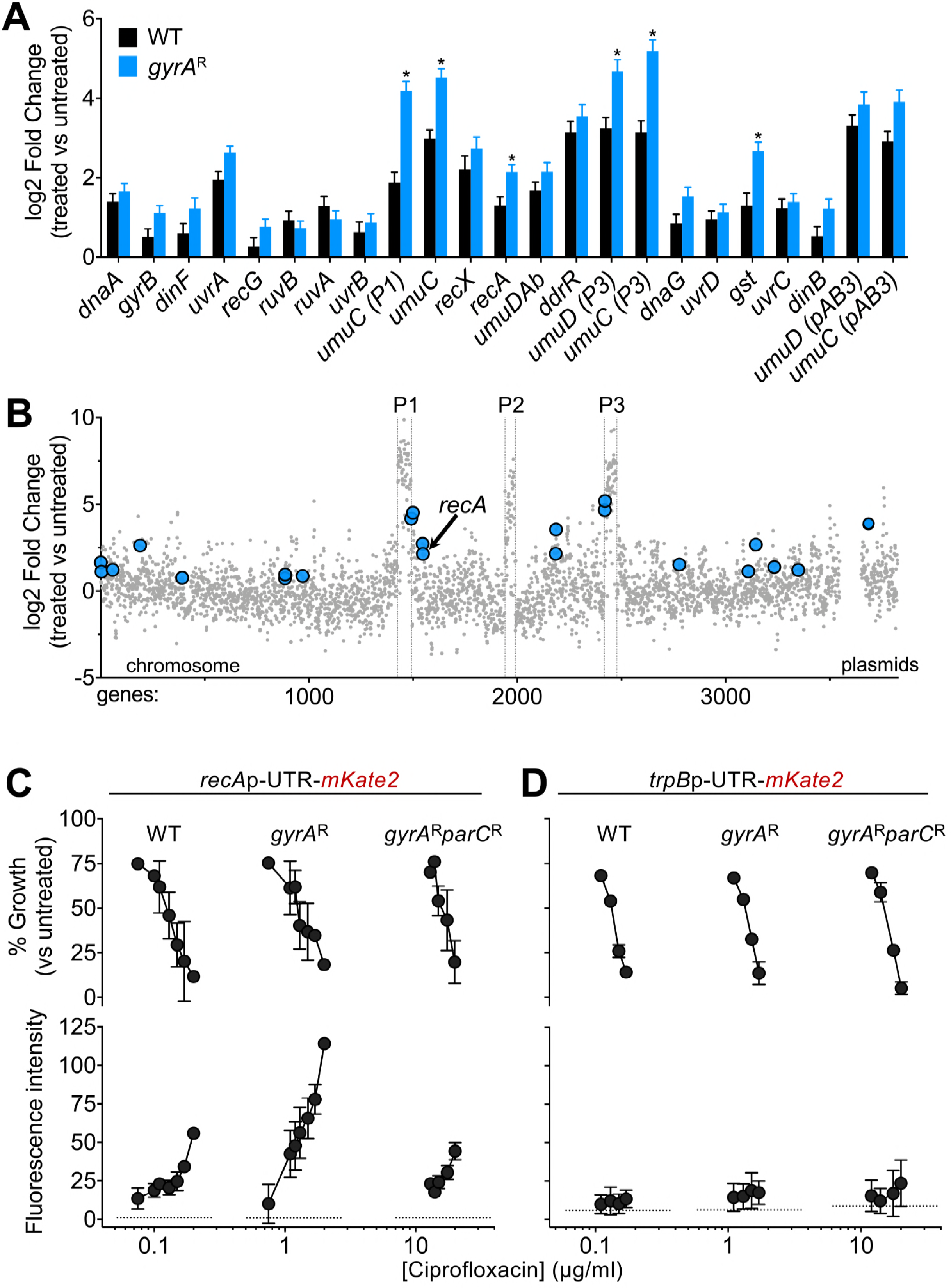
Enhanced SOS response induction in *gyrA*^R^-single mutants exposed to ciprofloxacin. (A-B) SOS response genes are induced during growth with ciprofloxacin. (A) RNA-seq data reveal DNA damage/SOS response induction during growth with ciprofloxacin. Bars show log2 fold change ± SEM (n = 3) for WT or *gyrA*^R^-single mutant treated with ciprofloxacin concentrations resulting in ~70% growth inhibition (condition *ii* from Fig. 5). *, p<0.05, unpaired t test. (B) Location of DNA damage/SOS response genes induced in *gyrA*^R^-single mutant strain. x-axis indicates gene position along the *A. baumannii* chromosome, y-axis indicates the log2 fold change (Cip 1.1μg/ml vs untreated, *gyrA*^R^-single mutant) from previously presented RNA-seq data. (C-D) Fluorescence reporter assays demonstrate enhanced *recA* gene expression in *gyrA*^R^-single mutant. Strains of the indicated genotype harboring (C) pCC1 (mKate2 fusion to *recA* promoter plus 5’ untranslated region (UTR)) or (D) pCC7 (*trpB* promoter replacing *recA* promoter in pCC1) were cultured as in RNA-seq experiments. Growth inhibition relative to untreated control was calculated (top). Average mKate2 intensity per cell within each sample was measured by fluorescence microscopy, and median fluorescence values of the population were determined (bottom). Data points represent the average inhibition values (top) or average of median fluorescence values (bottom) ± SD from n ≥ 2 biological replicates pooled from multiple independent experiments. Dotted lines denote fluorescence intensity of untreated samples.

## Discussion

In this study we exploited the dual-target nature of fluoroquinolone antibiotics to uncover how resistance alleles acquired in target enzymes modulate the landscape of intrinsic resistance. Using Tn-seq, we performed comprehensive screens for determinants of resistance to the fluoroquinolone drug ciprofloxacin in isogeneic *A. baumannii* strains in which the drug preferentially targets either DNA gyrase or Topo IV. We found that the spectrum of genes contributing to intrinsic resistance was similar in genetic backgrounds in which both enzymes were WT or in which both enzymes had lowered drug sensitivity due to well-known acquired point mutations. Intrinsic resistance determinants identified in both backgrounds included the AdeIJK and AbeM efflux pumps, multiple subunits of the DNA recombination and repair machinery, a periplasmic protease CtpA, the cell wall transpeptidase PBP1A, and several proteins of unknown function. By contrast, interaction of ciprofloxacin with the *gyrA*^R^ single mutant in which Topo IV is the sensitive target dramatically altered the profile of genes that influence relative Tn-seq fitness. This altered fitness profile in *gyrA*^R^*parC*^+^ bacteria was shown to directly reflect amplification of DNA in the vicinity of two endogenous prophages due to preferential poisoning of Topo IV by ciprofloxacin. Prophage transcripts and the SOS pathway were also hyperactivated as a consequence of this drug-genotype interaction, likely facilitating the initiation of synthesis of prophage DNA in the *gyrA*^R^ strain.

Our data can be explained by the model shown in Fig. 7, if we assume that re-activation of *A. baumannii* prophages 1 and 3 requires the function of host DNA gyrase. In WT and the *gyrA*^R^ *parC*^R^ double mutant, DNA gyrase is the effective target blocked by ciprofloxacin at growth-inhibitory, sub-MIC drug concentrations. Prophage DNA synthesis is blocked despite induction of the DNA damage response and prophage gene transcription because, as postulated by the model, efficient replication of the prophage genomes requires functional host gyrase (Fig. 7A and C). By contrast, in *gyrA*^R^ bacteria, Topo IV is the preferred target of intoxication by ciprofloxacin. This interaction causes DNA lesions that robustly stimulate the SOS pathway and prophage transcription; further, host DNA gyrase is available to facilitate prophage genome replication because the GyrA S81L (*gyrA*^R^) variant is resistant to the intermediate concentrations of ciprofloxacin required for Topo IV poisoning (Fig. 7B). Therefore, the presence of the single *gyrA*^R^ resistance generates a liability that is not observed in other strains, resulting from the induction of potentially lethal prophage replication in the presence of fluoroquinolones.

**Fig. 7.**
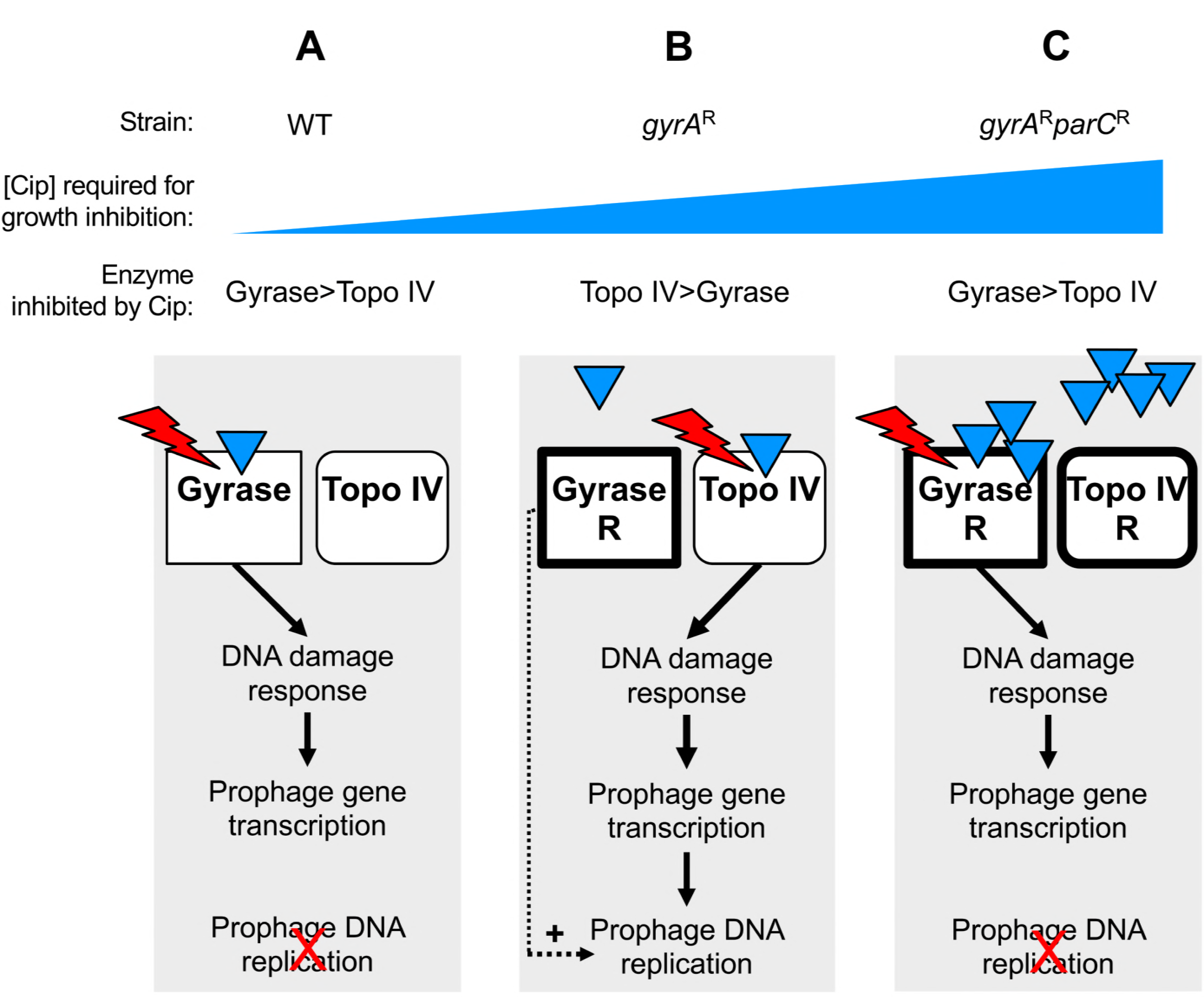
Model for resistance allele-dependent prophage amplification in *A. baumannii* exposed to sub-MIC fluoroquinolone stress. The model posits that prophage DNA replication depends on host DNA gyrase activity. (A) In WT cells, both gyrase (GyrA) and topo IV (ParC) are drug sensitive. Gyrase, which has higher affinity for CIP (blue triangles), is effectively targeted by the drug. Ciprofloxacin-corrupted gyrase results in double-strand DNA breaks that signal derepression of prophage gene expression. Prophage DNA replication cannot proceed, however, because gyrase function is blocked. (B) In single *gyrA*^R^ mutant cells, topo IV/ParC has higher affinity for ciprofloxacin than the resistant gyrase and is the effective drug target. Topo IV corruption results in a robust DNA damage response and activation of prophage gene expression, and gyrase-dependent prophage replication (prophages 1 and 3) proceeds because GyrA is not drug-inhibited. (C) In double *gyrA*^R^*parC*^R^ mutant cells growing at high drug concentrations, GyrA again has relatively higher affinity for ciprofloxacin than ParC and becomes the effective target despite the S81L drug binding site alteration. The resulting DNA lesions induce the SOS response and prophage gene expression, but prophage replication does not proceed efficiently because gyrase function is again blocked.

The central assumption of the model is plausible based on analogy with several other bacteriophage systems that have been shown to require host DNA gyrase for replication. Gyrase inhibitors (quinolones or aminocoumarins) inhibit phage DNA replication during lytic growth after infection (46-51), and disrupt induction of replicative transposition in Mu lysogens (52). Moreover, host gyrase is required for propagation of replication forks within supercoiled DNA substrates in reconstituted systems modeling phage lambda replication (53, 54). The importance of this enzyme class for replication of *A. baumannii* strain 17978 prophages is emphasized by the fact that they do not encode type II topoisomerases which are often encoded by bacteriophages to bypass a requirement for the host enzymes (55, 56).

An alternative model is that at the sub-MIC drug doses resulting in equivalent growth inhibition, gyrase poisoning results in DNA lesions that do not stimulate the SOS response above the threshold required for efficient prophage induction, in contrast to lesions caused by Topo IV poisoning. Arguing against this model are the observations that transcription of the SOS response gene *recA* and genes from all three prophages are strongly induced (25-50 fold, Figs. 5B and 6C) above baseline in WT cells, and Tn-seq fitness results showing the relatively similar importance of DNA damage repair enzymes across all strain backgrounds (Data Sets S1-S3). We showed that Topo IV intoxication stimulated the SOS pathway to a greater extent than that caused by gyrase poisoning, potentially contributing to the robust activation of prophages. This is consistent with previous findings that Topo IV and gyrase intoxication can be distinguished by several characteristics. Topo IV lesions result in slower inhibition of DNA synthesis and are thought to be more readily reversed by recombinational repair, resulting in lower cytotoxicity at given drug concentrations (16). These less toxic lesions could potentially expose more numerous or potent signals for the SOS response that could result in prophage induction.

In contrast with induction of prophages 1 and 3, ciprofloxacin-induced DNA replication was not observed with prophage 2 despite activation of prophage gene transcription. One possible explanation is that prophage 2 is defective for DNA replication. We consider this unlikely because transposon insertions were unobtainable in a phage locus (ACX60_RS10145) encoding a putative Cro/Cl family repressor (Data Sets S1-S3), indicating that this prophage has the potential for lytic replication in the absence of a protein controlling lysogeny maintenance. Consistent with the potential of all three prophages (including prophage 2) for replication, mobilized DNA corresponding to each of the three prophages was detected in phage particles resulting from treatment of WT *A. baumannii* 17978 with mitomycin C, which damages DNA directly without dependence on interactions with DNA topoisomerases (44). An alternative explanation for the lack of prophage 2 DNA amplification observed with fluoroquinolone treatment in our study is that its replication depends on both DNA gyrase and Topo IV.

The findings described here have implications for the evolution of antibiotic resistance in *A. baumannii* and other Gram-negative organisms. They indicate that in the trajectory toward high-level fluoroquinolone resistance, intermediate states with moderate-level fluoroquinolone resistance (exemplified by the *gyrA*^R^ single mutant) are those that possess highest potential for prophage induction during growth with continued drug exposure. Depending on the outcome of phage-host interactions, prophage hyperamplification within these bacteria could impose a fitness burden or could result in cell death if productive lysis ensues, representing additional selective pressures to influence evolution when under stress from the inducing antibiotic. Acquiring the subsequent *parC* mutation would answer this pressure and result in high-level fluoroquinolone resistance. It is notable that hyperamplified and hyperexpressed DNA within or adjacent to induced prophages include multiple *umuCD* paralogs encoding mutagenic DNA polymerases (57), whose higher levels of activity could increase mutation frequency and hasten bacterial adaptation in drug-treated *gyrA*^R^ single mutants. Moreover, these findings raise the possibility of enhanced horizontal transfer of phage-encoded and phage-proximal genes as a consequence of fluoroquinolone-*gyrA*^R^ interactions. The relationship between stepwise fluoroquinolone resistance and induction of prophages by this drug class may play out differently with Gram-positive organisms in which Topo IV is typically the sensitive initial target as opposed to gyrase (6). If fluoroquinolone-prophage dynamics in such bacteria have features that accord with the model proposed here, WT strains with two sensitive *parC* and *gyrA* alleles may represent the state with higher potential for drug-induced prophage replication than derivatives that have acquired single-step target-resistance mutations.

In summary, we have demonstrated that in the course of stepwise selection for high drug resistance, intermediate steps result in unexpected nodes of hypersensitivity that place both added pressure for acquisition of additional drug resistant alleles, as well as inducing the enzymatic machinery that drives acquisition of drug resistance. Future work on analysis of proteins that modulate the survival of drug resistant mutants should uncover strategies that allow these variants to be targeted therapeutically.

## Materials and Methods

### Bacterial strains, growth conditions, and antibiotics

Bacterial strains used in this work are described in Table S1. *A. baumannii* strains were derivatives of ATCC 17978. Bacterial cultures were grown at 37°C in Lysogeny Broth (LB) (10 g/L tryptone, 5 g/L yeast extract, 10 g/L NaCl) in flasks with shaking or in tubes on a roller drum. Growth was monitored by measuring absorbance at 600nm via a spectrophotometer. LB agar was supplemented with antibiotics [ampicillin (Amp, 50-100 μg/ml), carbenicillin (Cb, 50-100 μg/ml), kanamycin (Km, 10-20 μg/ml), mecillinam, ciprofloxacin] or sucrose as needed (Sigma Aldrich).

### Molecular cloning and mutant construction

Oligonucleotide primers and plasmids used in this study are listed in Table S2. Single *gyrA*S81L (*gyrA*^R^) and *parC*S84L (*parC* ^R^) point mutations were generated by cloning the respective genomic fragments in pUC18, followed by inverse PCR and self-ligation or amplification and substitution of a mutated gene fragment. In-frame deletions of ciprofloxacin-resistance genes were generated as described (58). Constructs were subcloned in pSR47S and used to isolate *A. baumannii* mutants via homologous recombination with two selection steps (58). The *gyrA*^R^ *parC* ^R^ double mutant was isolated by selection of a derivative of the *gyrA*^R^ strain able to grow on LB agar containing 8 μg/ml ciprofloxacin. A *pbp1a* mutant (N178TfsX27) was isolated as a derivative of ATCC 17978 selected on LB agar containing 64 μg/ml mecillinam.

### Antibiotic susceptibility assays

For growth curve analysis, cultures were seeded at A_600_ = 0.003 in 100μl of broth in wells of a 96-well microtiter plate and growth monitored during incubation at 37°C with orbital shaking in a Tecan M200 Pro plate reader. MIC tests were performed under the conditions above using serial 2-fold dilutions of drug; the MIC was the lowest drug concentration preventing growth above A_600_ = 0.05 after 16 hours.

### Construction of transposon mutant libraries

Plasmid pDL1073 was employed for transposon mutagenesis. pDL1073 contains a Km^R^ Tn10 derivative, an altered target-specificity Tn10 transposase gene downstream of the phage lambda P_L_ promoter, a pSC101ts origin of replication, and β-lactamase (Amp^R^; Fig. S1). pDL1073 does not replicate in *A. baumannii* at 37°C allowing efficient detection at this temperature of transposition after delivery via electroporation. *A. baumannii* cells (50μl) were combined with 100ng pDL1073 and electroporated via a BioRad Gene Pulser (0.1cm gap length cuvette; 200Ω, 25μF, and 1.8kV). Electroporated cells were diluted with SOC broth and immediately spread onto membrane filters (0.45μm pore size) overlaid on pre-warmed LB agar plates. After incubating 2 hours at 37°C, the filter membranes were transferred to pre-warmed LB agar plates containing 20μg/ml Km and incubated at 37°C overnight to select for transposon mutants. Bacterial colonies were lifted from the filter by agitation in sterile PBS. Glycerol was added to 10% (v/v), and pooled mutant suspensions were aliquoted and stored at -80°C. 11-15 independent pools each consisting of approximately 6,000-18,000 mutants were generated in each strain background.

### Tn-seq fitness measurements

Transposon library aliquots were thawed, vortexed, diluted to A_600_ = 0.1 and grown to A_600_ = 0.2 in LB. Cultures were then back-diluted to A_600_ = 0.003 in 10ml LB without drug or with graded concentrations of ciprofloxacin. Parallel cultures were grown at 37°C for approximately 8 generations to A_600_ = 0.5-1. Samples taken at the start (t_1_) and end (t_2_) of this outgrowth were stored at -20°C. 11 to 15 independent transposon libraries were analyzed with each strain background. With WT libraries, treatments with 0.075 μg/ml and 0.09-0.1 μg/ml ciprofloxacin were performed in parallel with the same untreated control.

### Tn-seq Illumina library preparation

Genomic DNA was extracted from t_1_ and t_2_ samples (Qiagen DNeasy Kit) and quantified by a SYBR green microtiter assay. Transposon-adjacent DNA was amplified for Illumina sequencing using a modification of the Nextera^TM^ DNA Library Prep method (Illumina). 30ng of genomic DNA was used as input in a 10μl tagmentation reaction. Reaction conditions were 55°C for 5min followed by inactivation at 95°C for 0.5min. Transposon-adjacent genomic DNA was amplified by adding 40μl of PCR master mix containing primers olj928 and Nextera 2A-R (0.6μM final) and Q5 High-Fidelity polymerase (NEB). Reaction conditions were 98°C for 10s, 65°C for 20s, and 72°C for 1min (30 cycles), followed by a final extension at 72°C for 2min. A second PCR was performed using nested, indexed primers. This reaction contained 0.5μL of the first PCR reaction, Left Tn10 indexing primer (0.6μM), Right indexing primer (0.6μM) and Q5 polymerase in a 50μl final volume. Reaction conditions were 98°C for 10s, 65°C for 20s, and 72°C for 1min (12 cycles of), followed by a final extension at 72°C for 2min. A sample of the second PCR product was imaged after separation on a 2% agarose/TAE gel containing SYBR Safe dye. Samples were multiplexed based on signal intensity in the 250-600bp region and purified (Qiagen QIAquick). 15-20pmol of DNA was used as template in a 50μl reconditioning reaction containing adapter-specific primers P1 and P2 (0.6μM) and Q5 polymerase. Reaction conditions were 95°C for 1min, 0.1°C/sec ramp to 64°C, 64°C for 20s, 72°C for 10min. Samples were purified (Qiagen QIAquick), followed by quantification and size selection (250-600bp, Pippin HT) by the Tufts University Genomics Core Facility (TUCF-Genomics). Libraries were sequenced (single-end 50bp) using custom primer olk115 on a HiSeq2500 with High Output V4 chemistry at TUCF-Genomics.

### Tn-seq data analysis

Reads were demultiplexed, quality-filtered and clipped of adapters before serving as input for mapping and fitness calculations (31). Reads were mapped to the *A. baumannii* 17978-mff chromosome (NZ_CP012004) and plasmids (NC_009083, NC_009084, and NZ_CP012005) using previously described parameters (59). Fitness values for each transposon mutant were calculated by comparing mutant vs population-wide expansion between t_1_ and t_2_ (31). Per-gene average fitness and SD were then computed from fitness scores for all insertion mutations within a gene across multiple parallel transposon pools. Differences in average gene fitness between treated and untreated conditions (W_diff_) were considered significant if they fulfilled the following 3 criteria, with minor modification from those previously described (33): per-gene fitness must be calculated from at least 3 data points, the magnitude of W_diff_ must be > 10%, and q value must be < 0.05 in an unpaired t-test with FDR controlled by the 2-stage step-up method of Benjamini, Krieger and Yekutieli (GraphPad Prism 7). Per-insertion fitness scores within a given genomic region were visualized using Integrative Genomics Viewer software (60) after aggregating all scores across multiple independent transposon mutant libraries using the SingleFitness Perl script (61).

### Whole-genome sequencing of individual strains subjected to ciprofloxacin

WT, *gyrA*^R^, or *gyrA*^R^ *parC*^R^ strains were grown from single colonies to early post-exponential phase and back-diluted to A_600_ 0.003. Parallel cultures were grown for 2.5 hours in the absence of treatment, or 3.5 hours in the presence of ciprofloxacin treatment. DNA was extracted (Qiagen DNeasy) and Illumina sequencing libraries were amplified and sequenced as described (34). After mapping to NZ_CP012004, coverage files were generated from the resulting BAM files using deepTools, with reads normalized to counts per million (62).

### Transcriptional profiling

Cultures were diluted to A_600_ 0.003 and grown for 2.5 hours (untreated) or 3.5 hours (ciprofloxacin treated). Cultures were mixed with an equal volume of ice-cold acetone:ethanol (1:1) and stored at -80°C. Cells were thawed and washed with TE and RNA was extracted (Qiagen RNeasy). RNA samples were diluted, combined with SUPERase-in (Invitrogen), and processed via the RNAtag-seq method (42). Illumina cDNA sequencing libraries were sequenced and reads processed as described (63). Differential expression was calculated using DESeq2 (64).

### Fluorescence reporter assays

Strains containing pCC1 or pCC7 were cultured in the presence or absence of ciprofloxacin as in RNA-seq experiments. Cells were immobilized on agarose pads and imaged on a Leica AF6000 microscope using a 100X/1.3 objective and TX2 filtercube (excitation: BP 560/40, dichromatic mirror 595, emission: BP 645/75). MicrobeJ (65) was used to measure background-corrected mean fluorescence intensity per cell. Median cellular fluorescence intensities from populations of at least 100 bacteria were determined, and median values across multiple independent experiments were averaged.

### Accession Number(s)

Sequencing reads analyzed in this study were deposited into SRA database as: SRP157243 (Tn-seq), PRJNA495614 (RNA-seq), and PRJNA495623 (Whole genome sequencing).

## Acknowledgements

This work was supported by NIAID awards U01AI124302 to RRI and TVO, R21AI128328 to RRI, and F32AI098358 to EG. NSF REU Site Award #1757443 to the Northeastern University Department of Biology supported the research of ELW. We thank Carly Ching and Veronica Godoy for gift of pCC1 and pCC7, and Amy Tang for assistance with bioinformatics analysis.

## Supplemental Figure Legends

**Fig. S1. pDL1073 feature map.**

**Fig. S2. Sub-MIC ciprofloxacin treatment of *A. baumannii* with *gyrA*^WT^ and *parC*^WT^ alleles does not significantly alter Tn-seq fitness values assigned to prophage region genes.** (A) The Tn-seq dataset shown in Fig. 1B (WT background +/-treatment with ciprofloxacin 0.09-0.1 μg/ml) was reanalyzed to highlight fitness values associated with genes within prophage regions P1-P3. Prophages regions are highlighted with color indicated in the key.

**Fig. S3. Comparison of transcription levels between ciprofloxacin-treated cultures of WT and *gyrA*^R^ reveals enhanced prophage gene expression in the *gyrA*^R^ single mutant.** Strains were grown in the absence or presence of ciprofloxacin at concentrations shown in Fig. 5A, and RNA-seq data were analyzed such that WT and *gyrA*^R^ strains were directly compared at each condition. Plots show log2 fold change (WT vs *gyrA*^R^) of each gene in order of position on the chromosome or plasmids (pAB1-3).

**Fig. S4. Fluorescence microscopy analysis of SOS response in individual cells subjected to ciprofloxacin**. Strains harboring *recA-mKate2* were cultured and analyzed by fluorescence microscopy as described in legend to Fig. 6C. (A) Phase contrast (rows 1 and 3) and fluorescence (rows 2 and 4) microscopy images from one representative experiment used to quantify *recA-mKate2* signal in Fig. 6C. Cells of the indicated genotype were treated with the noted concentration of ciprofloxacin resulting in similar degrees of growth inhibition (see panel C, bottom three growth inhibition data points). (B) Population fluorescence analysis from one representative experiment contributing to Fig. 6C quantifying SOS response to increasing ciprofloxacin dose in different strain backgrounds harboring *recA-mKate2*. In the same experiment shown in panel A, 4 different ciprofloxacin concentrations were tested per strain. Average mKate2 intensity per cell was measured by fluorescence microscopy. Each data point represents average fluorescence intensity of a single cell (at least 100 cells per condition were analyzed). Bars indicate median values. (C) Growth inhibition relative to untreated control resulting from the ciprofloxacin exposures in the representative experiment shown in panel B.

**Table S1.** Bacterial strains and plasmids used in this study.

**Table S2.** Oligonucleotide primers used in this study.

**Data Set S1.** Tn-seq fitness data - WT.

**Data Set S2.** Tn-seq fitness data - *gyrA*^R^.

**Data Set S3.** Tn-seq fitness data - *gyrA*^R^ *parC*^R^.

**Data Set S4.** RNA-seq data.

## References

1. Tacconelli E, Carrara E, Savoldi A, Harbarth S, Mendelson M, Monnet DL, Pulcini C, Kahlmeter G, Kluytmans J, Carmeli Y, Ouellette M, Outterson K, Patel J, Cavaleri M, Cox EM, Houchens CR, Grayson ML, Hansen P, Singh N, Theuretzbacher U, Magrini N, Group WHOPPLW. 2018. Discovery, research, and development of new antibiotics: the WHO priority list of antibiotic-resistant bacteria and tuberculosis. Lancet Infect Dis 18:318–327.

2. Warner WA, Kuang SN, Hernandez R, Chong MC, Ewing PJ, Fleischer J, Meng J, Chu S, Terashita D, English L, Chen W, Xu HH. 2016. Molecular characterization and antimicrobial susceptibility of Acinetobacter baumannii isolates obtained from two hospital outbreaks in Los Angeles County, California, USA. BMC Infect Dis 16:194

3. Kim D, Ahn JY, Lee CH, Jang SJ, Lee H, Yong D, Jeong SH, Lee K. 2017. Increasing Resistance to Extended-Spectrum Cephalosporins, Fluoroquinolone, and Carbapenem in Gram-Negative Bacilli and the Emergence of Carbapenem Non-Susceptibility in Klebsiella pneumoniae: Analysis of Korean Antimicrobial Resistance Monitoring System (KARMS) Data From 2013 to 2015. Ann Lab Med 37:231–239.

4. Blanco N, Harris AD, Rock C, Johnson JK, Pineles L, Bonomo RA, Srinivasan A, Pettigrew MM, Thom KA, the CDCEP. 2018. Risk Factors and Outcomes Associated with Multidrug-Resistant Acinetobacter baumannii upon Intensive Care Unit Admission. Antimicrob Agents Chemother 62.

5. Drlica K, Hiasa H, Kerns R, Malik M, Mustaev A, Zhao X. 2009. Quinolones: action and resistance updated. Curr Top Med Chem 9:981–98.

6. Hooper DC. 1999. Mechanisms of fluoroquinolone resistance. Drug Resist Updat 2:38–55.

7. Jacoby GA. 2005. Mechanisms of resistance to quinolones. Clin Infect Dis 41 Suppl 2:S120–6.

8. Hujer KM, Hujer AM, Hulten EA, Bajaksouzian S, Adams JM, Donskey CJ, Ecker DJ, Massire C, Eshoo MW, Sampath R, Thomson JM, Rather PN, Craft DW, Fishbain JT, Ewell AJ, Jacobs MR, Paterson DL, Bonomo RA. 2006. Analysis of antibiotic resistance genes in multidrug-resistant Acinetobacter sp. isolates from military and civilian patients treated at the Walter Reed Army Medical Center. Antimicrob Agents Chemother 50:4114–23.

9. Valentine SC, Contreras D, Tan S, Real LJ, Chu S, Xu HH. 2008. Phenotypic and molecular characterization of Acinetobacter baumannii clinical isolates from nosocomial outbreaks in Los Angeles County, California. J Clin Microbiol 46:2499–507.

10. Fernando D, Zhanel G, Kumar A. 2013. Antibiotic resistance and expression of resistance-nodulation-division pump- and outer membrane porin-encoding genes in Acinetobacter species isolated from Canadian hospitals. Can J Infect Dis Med Microbiol 24:17–21.

11. Rumbo C, Gato E, Lopez M, Ruiz de Alegria C, Fernandez-Cuenca F, Martinez-Martinez L, Vila J, Pachon J, Cisneros JM, Rodriguez-Bano J, Pascual A, Bou G, Tomas M, Spanish Group of Nosocomial I, Mechanisms of A, Resistance to A, Spanish Society of Clinical M, Infectious D, Spanish Network for Research in Infectious D. 2013. Contribution of efflux pumps, porins, and beta-lactamases to multidrug resistance in clinical isolates of Acinetobacter baumannii. Antimicrob Agents Chemother 57:5247–57.

12. Yoon EJ, Chabane YN, Goussard S, Snesrud E, Courvalin P, De E, Grillot-Courvalin C. 2015. Contribution of resistance-nodulation-cell division efflux systems to antibiotic resistance and biofilm formation in Acinetobacter baumannii. MBio 6:e00309–15.

13. Poole K. 2000. Efflux-mediated resistance to fluoroquinolones in gram-negative bacteria. Antimicrob Agents Chemother 44:2233–41.

14. Recacha E, Machuca J, Diaz de Alba P, Ramos-Guelfo M, Docobo-Perez F, Rodriguez-Beltran J, Blazquez J, Pascual A, Rodriguez-Martinez JM. 2017. Quinolone Resistance Reversion by Targeting the SOS Response. MBio 8.

15. Cirz RT, Chin JK, Andes DR, de Crecy-Lagard V, Craig WA, Romesberg FE. 2005. Inhibition of mutation and combating the evolution of antibiotic resistance. PLoS Biol 3:e176

16. Khodursky AB, Cozzarelli NR. 1998. The mechanism of inhibition of topoisomerase IV by quinolone antibacterials. J Biol Chem 273:27668–77.

17. Mo CY, Manning SA, Roggiani M, Culyba MJ, Samuels AN, Sniegowski PD, Goulian M, Kohli RM. 2016. Systematically Altering Bacterial SOS Activity under Stress Reveals Therapeutic Strategies for Potentiating Antibiotics. mSphere 1.

18. Nichols RJ, Sen S, Choo YJ, Beltrao P, Zietek M, Chaba R, Lee S, Kazmierczak KM, Lee KJ, Wong A, Shales M, Lovett S, Winkler ME, Krogan NJ, Typas A, Gross CA. 2011. Phenotypic landscape of a bacterial cell. Cell 144:143–56.

19. Sutherland JH, Tse-Dinh YC. 2010. Analysis of RuvABC and RecG involvement in the escherichia coli response to the covalent topoisomerase-DNA complex. J Bacteriol 192:4445–51.

20. Tamae C, Liu A, Kim K, Sitz D, Hong J, Becket E, Bui A, Solaimani P, Tran KP, Yang H, Miller JH. 2008. Determination of antibiotic hypersensitivity among 4,000 single-gene-knockout mutants of Escherichia coli. J Bacteriol 190:5981–8.

21. Urios A, Herrera G, Aleixandre V, Blanco M. 1990. Expression of the recA gene is reduced in Escherichia coli topoisomerase I mutants. Mutat Res 243:267–72.

22. Brazas MD, Breidenstein EB, Overhage J, Hancock RE. 2007. Role of lon, an ATP-dependent protease homolog, in resistance of Pseudomonas aeruginosa to ciprofloxacin. Antimicrob Agents Chemother 51:4276–83.

23. Breidenstein EB, Khaira BK, Wiegand I, Overhage J, Hancock RE. 2008. Complex ciprofloxacin resistome revealed by screening a Pseudomonas aeruginosa mutant library for altered susceptibility. Antimicrob Agents Chemother 52:4486–91.

24. van Opijnen T, Camilli A. 2012. A fine scale phenotype-genotype virulence map of a bacterial pathogen. Genome Res 22:2541–51.

25. Kelley WL. 2006. Lex marks the spot: the virulent side of SOS and a closer look at the LexA regulon. Mol Microbiol 62:1228–38.

26. Robinson A, Brzoska AJ, Turner KM, Withers R, Harry EJ, Lewis PJ, Dixon NE. 2010. Essential biological processes of an emerging pathogen: DNA replication, transcription, and cell division in Acinetobacter spp. Microbiol Mol Biol Rev 74:273–97.

27. Macguire AE, Ching MC, Diamond BH, Kazakov A, Novichkov P, Godoy VG. 2014. Activation of phenotypic subpopulations in response to ciprofloxacin treatment in Acinetobacter baumannii. Mol Microbiol 92:138–52.

28. Aranda J, Bardina C, Beceiro A, Rumbo S, Cabral MP, Barbe J, Bou G. 2011. Acinetobacter baumannii RecA protein in repair of DNA damage, antimicrobial resistance, general stress response, and virulence. J Bacteriol 193:3740–7.

29. Saroj SD, Clemmer KM, Bonomo RA, Rather PN. 2012. Novel mechanism for fluoroquinolone resistance in Acinetobacter baumannii. Antimicrob Agents Chemother 56:4955–7.

30. Gallagher LA, Lee SA, Manoil C. 2017. Importance of Core Genome Functions for an Extreme Antibiotic Resistance Trait. MBio 8.

31. van Opijnen T, Camilli A. 2013. Transposon insertion sequencing: a new tool for systems-level analysis of microorganisms. Nat Rev Microbiol 11:435–42.

32. van Opijnen T, Bodi KL, Camilli A. 2009. Tn-seq: high-throughput parallel sequencing for fitness and genetic interaction studies in microorganisms. Nat Methods 6:767–72.

33. van Opijnen T, Dedrick S, Bento J. 2016. Strain Dependent Genetic Networks for Antibiotic-Sensitivity in a Bacterial Pathogen with a Large Pan-Genome. PLoS Pathog 12:e1005869

34. Geisinger E, Mortman NJ, Vargas-Cuebas G, Tai AK, Isberg RR. 2018. A global regulatory system links virulence and antibiotic resistance to envelope homeostasis in Acinetobacter baumannii. PLoS Pathog 14:e1007030

35. Drlica K, Zhao X. 1997. DNA gyrase, topoisomerase IV, and the 4-quinolones. Microbiol Mol Biol Rev 61:377–92.

36. Su XZ, Chen J, Mizushima T, Kuroda T, Tsuchiya T. 2005. AbeM, an H+-coupled Acinetobacter baumannii multidrug efflux pump belonging to the MATE family of transporters. Antimicrob Agents Chemother 49:4362–4.

37. Knauf GA, Cunningham AL, Kazi MI, Riddington IM, Crofts AA, Cattoir V, Trent MS, Davies BW. 2018. Exploring the Antimicrobial Action of Quaternary Amines against Acinetobacter baumannii. MBio 9.

38. Gomez JE, Kaufmann-Malaga BB, Wivagg CN, Kim PB, Silvis MR, Renedo N, Ioerger TR, Ahmad R, Livny J, Fishbein S, Sacchettini JC, Carr SA, Hung DT. 2017. Ribosomal mutations promote the evolution of antibiotic resistance in a multidrug environment. Elife 6.

39. Bollenbach T, Quan S, Chait R, Kishony R. 2009. Nonoptimal microbial response to antibiotics underlies suppressive drug interactions. Cell 139:707–18.

40. Shippy DC, Fadl AA. 2015. RNA modification enzymes encoded by the gid operon: Implications in biology and virulence of bacteria. Microb Pathog 89:100–7.

41. Hujer KM, Hujer AM, Endimiani A, Thomson JM, Adams MD, Goglin K, Rather PN, Pennella TT, Massire C, Eshoo MW, Sampath R, Blyn LB, Ecker DJ, Bonomo RA. 2009. Rapid determination of quinolone resistance in Acinetobacter spp. J Clin Microbiol 47:1436–42.

42. Shishkin AA, Giannoukos G, Kucukural A, Ciulla D, Busby M, Surka C, Chen J, Bhattacharyya RP, Rudy RF, Patel MM, Novod N, Hung DT, Gnirke A, Garber M, Guttman M, Livny J. 2015. Simultaneous generation of many RNA-seq libraries in a single reaction. Nat Methods 12:323–5.

43. Simmons LA, Foti JJ, Cohen SE, Walker GC. 2008. The SOS Regulatory Network. EcoSal Plus 3.

44. Hare JM, Ferrell JC, Witkowski TA, Grice AN. 2014. Prophage induction and differential RecA and UmuDAb transcriptome regulation in the DNA damage responses of Acinetobacter baumannii and Acinetobacter baylyi. PLoS One 9:e93861

45. Ching C, Gozzi K, Heinemann B, Chai Y, Godoy VG. 2017. RNA-Mediated cis Regulation in Acinetobacter baumannii Modulates Stress-Induced Phenotypic Variation. J Bacteriol 199.

46. Erb ML, Kraemer JA, Coker JK, Chaikeeratisak V, Nonejuie P, Agard DA, Pogliano J. 2014. A bacteriophage tubulin harnesses dynamic instability to center DNA in infected cells. Elife 3.

47. Constantinou A, Voelkel-Meiman K, Sternglanz R, McCorquodale MM, McCorquodale DJ. 1986. Involvement of host DNA gyrase in growth of bacteriophage T5. J Virol 57:875–82.

48. Alonso JC, Sarachu AN, Grau O. 1981. DNA gyrase inhibitors block development of Bacillus subtilis bacteriophage SP01. J Virol 39:855–60.

49. Alcorlo M, Salas M, Hermoso JM. 2007. In vivo DNA binding of bacteriophage GA-1 protein p6. J Bacteriol 189:8024–33.

50. Kreuzer KN, Cozzarelli NR. 1979. Escherichia coli mutants thermosensitive for deoxyribonucleic acid gyrase subunit A: effects on deoxyribonucleic acid replication, transcription, and bacteriophage growth. J Bacteriol 140:424–35.

51. Itoh T, Tomizawa JI. 1977. Involvement of DNA gyrase in bacteriophage T7 DNA replication. Nature 270:78–80.

52. Sokolsky TD, Baker TA. 2003. DNA gyrase requirements distinguish the alternate pathways of Mu transposition. Mol Microbiol 47:397–409.

53. Dodson M, Echols H, Wickner S, Alfano C, Mensa-Wilmot K, Gomes B, LeBowitz J, Roberts JD, McMacken R. 1986. Specialized nucleoprotein structures at the origin of replication of bacteriophage lambda: localized unwinding of duplex DNA by a six-protein reaction. Proc Natl Acad Sci U S A 83:7638–42.

54. Mensa-Wilmot K, Seaby R, Alfano C, Wold MC, Gomes B, McMacken R. 1989. Reconstitution of a nine-protein system that initiates bacteriophage lambda DNA replication. J Biol Chem 264:2853–61.

55. Kreuzer KN. 1998. Bacteriophage T4, a model system for understanding the mechanism of type II topoisomerase inhibitors. Biochim Biophys Acta 1400:339–47.

56. Huang WM, Wei LS, Casjens S. 1985. Relationship between bacteriophage T4 and T6 DNA topoisomerases. T6 39-protein subunit is equivalent to the combined T4 39- and 60-protein subunits. J Biol Chem 260:8973–7.

57. Sutton MD, Smith BT, Godoy VG, Walker GC. 2000. The SOS response: recent insights into umuDC-dependent mutagenesis and DNA damage tolerance. Annu Rev Genet 34:479–497.

58. Geisinger E, Isberg RR. 2015. Antibiotic modulation of capsular exopolysaccharide and virulence in Acinetobacter baumannii. PLoS Pathog 11:e1004691

59. Carter R, Wolf J, van Opijnen T, Muller M, Obert C, Burnham C, Mann B, Li Y, Hayden RT, Pestina T, Persons D, Camilli A, Flynn PM, Tuomanen EI, Rosch JW. 2014. Genomic analyses of pneumococci from children with sickle cell disease expose host-specific bacterial adaptations and deficits in current interventions. Cell Host Microbe 15:587–599.

60. Robinson JT, Thorvaldsdottir H, Winckler W, Guttman M, Lander ES, Getz G, Mesirov JP. 2011. Integrative genomics viewer. Nat Biotechnol 29:24–6.

61. McCoy KM, Antonio ML, van Opijnen T. 2017. MAGenTA: a Galaxy implemented tool for complete Tn-Seq analysis and data visualization. Bioinformatics 33:2781–2783.

62. Ramirez F, Dundar F, Diehl S, Gruning BA, Manke T. 2014. deepTools: a flexible platform for exploring deep-sequencing data. Nucleic Acids Res 42:W187–91.

63. Jensen PA, Zhu Z, van Opijnen T. 2017. Antibiotics Disrupt Coordination between Transcriptional and Phenotypic Stress Responses in Pathogenic Bacteria. Cell Rep 20:1705–1716.

64. Love MI, Huber W, Anders S. 2014. Moderated estimation of fold change and dispersion for RNA-seq data with DESeq2. Genome Biol 15:550

65. Ducret A, Quardokus EM, Brun YV. 2016. MicrobeJ, a tool for high throughput bacterial cell detection and quantitative analysis. Nat Microbiol 1:16077

66. Arndt D, Grant JR, Marcu A, Sajed T, Pon A, Liang Y, Wishart DS. 2016. PHASTER: a better, faster version of the PHAST phage search tool. Nucleic Acids Res 44:W16–21.

